# High-throughput crystallographic fragment screening of Zika virus NS3 Helicase

**DOI:** 10.1101/2024.04.27.591279

**Authors:** Andre Schutzer Godoy, Charline Giroud, Nathalya C. M. R. Mesquita, Gabriela Dias Noske, Victor Oliveira Gawriljuk, Ryan M Lithgo, Blake H Balcomb, Jasmin Cara Aschenbrenner, Charles W.E. Tomlinson, Max Winokan, Jenke Scheen, Hugo MacDermott-Opeskin, Peter George Marples, Anu V. Chandran, Xiaomin Ni, Warren Thompson, Michael Fairhead, Daren Fearon, Lizbé Koekemoer, Mary-Ann Elvina Xavier, Martin Walsh, Glaucius Oliva, Frank von Delft

## Abstract

The Zika virus (ZIKV), discovered in Africa in 1947, swiftly spread across continents, causing significant concern due to its recent association with microcephaly in newborns and Guillain-Barré syndrome in adults. Despite a decrease in prevalence, the potential for a resurgence remains, necessitating urgent therapeutic interventions. Like other flaviviruses, ZIKV presents promising drug targets within its replication machinery, notably the NS3 helicase (NS3^Hel^) protein, which plays critical roles in viral replication. However, a lack of structural information impedes the development of specific inhibitors targeting NS3^Hel^. Here we applied high-throughput crystallographic fragment screening on ZIKV NS3^Hel^, which yielded structures that reveal 3D binding poses of 46 fragments at multiple sites of the protein, including 11 unique fragments in the RNA-cleft site. These fragment structures provide templates for direct design of hit compounds and should thus assist the development of novel direct-acting antivirals against ZIKV and related flaviviruses, thus opening a promising avenue for combating future outbreaks.

## Introduction

Since its discovery in Africa in 1947, the Zika virus (ZIKV) has traversed Asia and Polynesia before making its way to Brazil between May and December of 2013 (1, 2). From there, it swiftly spread across the globe (2). Initially, the symptoms of Zika fever were deemed too benign to raise significant concern within the medical community. However, this perception quickly shifted following the revelation of the correlation between ZIKV infection and the occurrence of microcephaly in newborns, as well as Guillain-Barré syndrome in adults (1, 3). Subsequently, additional neurological issues observed in newborns, including cortical malformations, ventriculomegaly, retinal damage, and arthrogryposis, were collectively termed Zika Congenital Syndrome (ZCS) (4–7). Although ZIKV has become less prevalent in recent times, cases continue to persist, raising concerns about the potential for a new pandemic outbreak. The absence of available vaccines or therapeutic agents for ZIKV underscores the urgent medical necessity, prompting ongoing efforts to identify potential drug candidates.

Such as Dengue virus (DENV) and Yellow Fever (YFV), the ZIKV belongs to the Flaviviridae family as a flavivirus (8). Its genome comprises a positive-sense, single-stranded RNA of approximately 10 kbp. (9). This is translated into a unique polyprotein which is proteolytically cleaved to originate 10 proteins separated in two groups: three structural proteins (Capsid (C), precursor-Membrane (prM/M) and Envelope (E)) and seven non-structural proteins - NSs - (NS1, NS2A, NS2B, NS3, NS4A NS4B and NS5) (9). Structural proteins form viral particles and nonstructural proteins participate in viral replication, virion assembly and evasion of the host immune response arboviruses (9). NS3 is composed by two domains: An N-terminal serine-protease (NS3^Pro^) and the C-terminal NTP-dependent RNA-helicase (NS3^Hel^). The NS3^Pro^ assemblies with the NS2B cofactor, forming the NS2B-NS3^Pro^, to cleave portions of the virion polyprotein after translation, whereas the NS3^Hel^ domain is responsible for double-stranded RNA-unwinding prior to RNA-polymerization (10). In addition to its enzymatic activity, NS3 engages in direct interactions with the viral largest membrane protein NS4B and NS5, leading to the formation of the viral replication complex (11, 12).

Considering the pivotal roles played by both enzymatic activities and interactions within the replication machinery in the process of viral replication, NS3 stands out as an exceptionally promising target for drug discovery (13, 14). Notably, currently ongoing clinical trial investigate a compound targeting NS3-NS4B interactions as a potential treatment for DENV (15, 16) Despite its pivotal role, the lack of structural information detailing specific interactions with NS3^Hel^ and small molecules poses a hindrance to the advancement of novel direct-acting antivirals aimed at targeting RNA-unwind activity. To address this gap, we employed high-throughput (HTP) crystallographic fragment screening to enable ZIKV NS3^Hel^ as a drug target, providing valuable insights for the development of a first-in-class RNA-unwind flaviviruses inhibitor.

## Results

### Overall fragment screening data processing

ZIKV NS3^Hel^ was successfully purified and crystallized. The initial ZIKV NS3^Hel^ crystals obtained were used to determine crystal structure with a good stereochemistry with 99.6 % of the residues in the most favored regions of the Ramachandran plot, after being refined to 1.9 Å with R_work_/R_free_ of 0.17/0.20 (PDB 6MH3). A single molecule was observed in the asymmetric unit, consistent with previous reported structures (17–19). Upon detailed inspection of crystal packing, significant solvent channels were observed providing access to essential sites within the protein structure. Soaking of crystals 20% DMSO for 2h showed minor effect on diffraction quality of the crystals, yielding datasets with resolution between 1.8 to 2.2 Å. Using these parameters, a total of 1,567 fragments were soaked from various library collections, resulting in 1,204 usable datasets. Impressively, 75% of these datasets diffracted above 2.5 Å, with an average resolution of 2.03 Å overall. PanDDA analysis revealed 451 events in distinct subsites of the protein. Following meticulous modeling, data curation, and cross-reviewing, a total of 45 ligand-bound and 1 ground-state structures of ZIKV NS3^Hel^ ranging from 1.36 to 2.1 Å resolution, were validated and subsequently deposited in the Protein Data Bank (PDB). PDB identifiers, names, ligand descriptors and data collection and refinement statistics are available in Supplementary Table 1.

### ZIKV NS3^Hel^ overall structure and fragments screening results

ZIKV NS3^Hel^ showcases a prototypical flavivirus RNA helicase insert, belonging to the helicase superfamily 2 (SF2), as evidenced by sequence comparisons and conserved motifs. It falls into the DEAH/RHA subfamilies, distinguished by the signature sequence within the Walker B motif (WB) (20–22). The structure of ZIKV NS3^Hel^ reveals a composition comprising approximately 16% β-sheets, 33% α-helices, and 51% coils, among other elements. These secondary structures delineate three distinct subdomains: Sub-domain I (DI), spanning residues 183-332; Sub-domain II (DII), encompassing residues 333-481, which exhibit characteristics akin to RecA-like domains; and Sub-domain III (DIII), spanning residues 482-617, presenting a Flavi-DEAD domain. The arrangement of these subdomains creates conserved clefts between them, referred to as RNA-cleft sites, which have been experimentally demonstrated to facilitate binding to single-strand RNAs. At the interface of DI and DII, another notably conserved feature is the presence of a cavity responsible for NTP-binding, which drives NTPase activity (10, 23).

Fragments were noted to be present throughout all surfaces and cavities of ZIKV NS3^Hel^ (Fig. 1a). Particularly noteworthy are three clusters that demonstrate promising characteristics and will be further discussed. To underscore the efficacy of HTP fragment screening, we conducted an alignment of 111 Flaviviridae PDB structures and visualized all small molecules present within them (Fig. 1b). The alignment reveals that much of the structural data available is redundant and primarily focuses on superficial ligands located at the NTP and RNA-entry sites (Fig. 1b). This comparison highlights how HTP fragment screening has successfully delved into chemical spaces that were previously unexplored for the SF2 helicase, shedding new light on the development of RNA binding competitors.

**Fig. 1.**
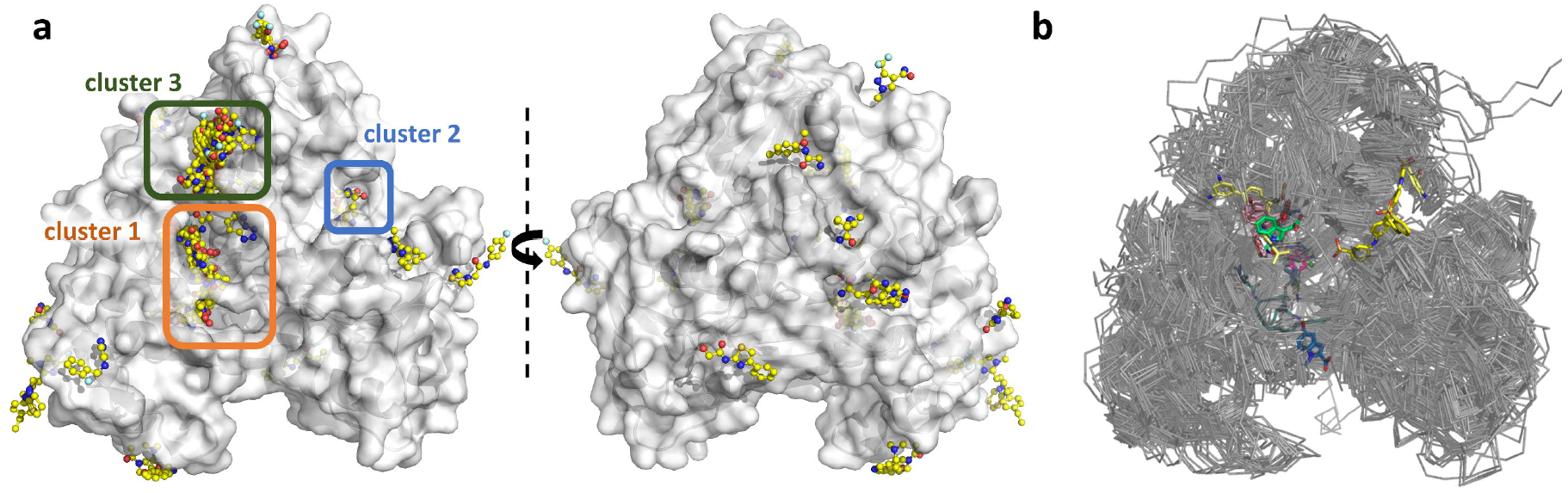
Fragment screening reveal multiple binding sites. (a) Surface view of ZIKV NS3^Hel^ fragment screening results in two orientations. Surface is colored in grey, and fragments are depicted as sticks with yellow carbons. Selected cluster of fragments are marked with orange (1), blue (2) and green (3) boxes. (b) Overlaid structures of 111 PBDs containing superfamily 2 helicases from *Flaviviridae* family. Small molecules are depicted as sticks.

### Fragments reveal new binding modes for the RNA-cleft site

In our HTP fragment screening initiative, we successfully elucidated the structure of ZIKV NS3^Hel^ bound to 11 fragments in the RNA-cleft site, marked as Cluster 1: Z235341991, Z1262246195, Z44584886, Z18618496, Z2678251369, POB0128, POB0008, Z3241250482, Z56772132, Z729726784 and Z203039992 (Fig 2a). These complexes are cataloged under the PDB codes 5RHG, 7G9K, 7G9M, 7G9T, 7G9Y, 7GA1, 7GA2, 7GA3, 8UM3, 8V7R, and 8V7U, respectively. Detailed information including PDB identifiers, fragment names, ligand descriptors, and data collection/refinement statistics can be found in Supplementary Table 1.

**Fig. 2.**
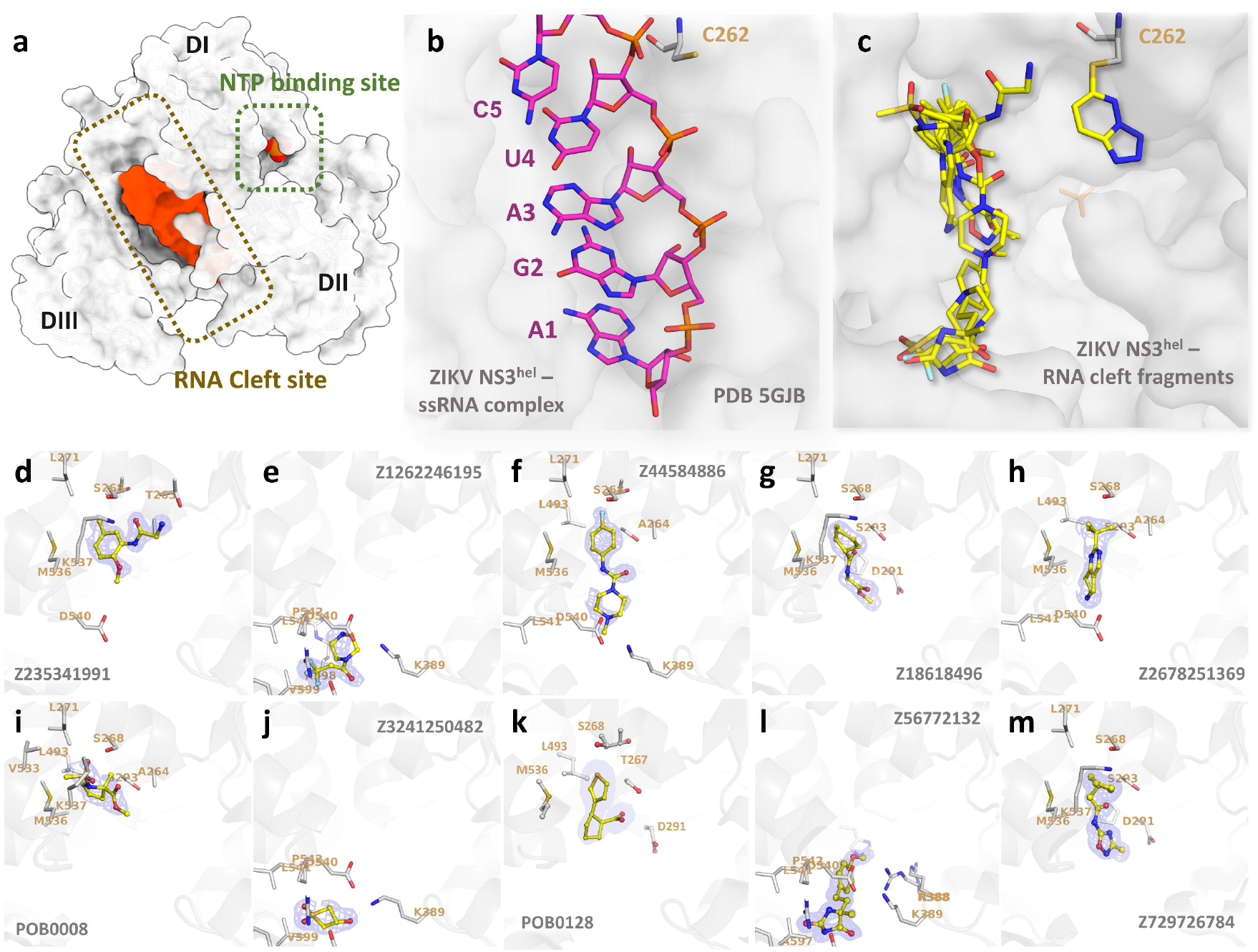
Overview of RNA cleft fragments obtained. (a) Surface view of the ZIKV NS3^Hel^ (depicted in grey) reveals distinct features: the NTP and RNA clefts are highlighted with green and brown boxes, respectively. Additionally, the RNA and phosphate surfaces are depicted in orange to accentuate the RNA binding cleft. (b) Detailed view of ZIKV Helicase from PDB 5GJB in complex with RNA. RNA is depicted as sticks. (c) ZIKV Helicase in complex with multiple fragments obtained for the RNA-cleft binding site. Fragments are depicted as sticks. Bellow, detailed view of structures with electron densities and structural details of fragments Z235341991 (d), Z1262246195 (e), Z44584886 (f), Z18618496 (g), Z2678251369 (h), POB0008 (i), Z3241250482 (j), POB0128 (k), Z56772132 (l) and Z729726784 (m). Fragments are show as sticks with yellow carbons, while protein interacting residues are depicted as grey sticks carbon. 2Fo-Fc map is depicted as blue mesh, showed here with σ = 1.0.

The alignment of all fragments from Cluster 1 elucidates their significant overlap with the RNA-bound structure from PDB 5GJB, particularly in the region responsible for base recognition (Fig 2b and c). Z235341991 is a N-(2-methoxy-5-methylphenyl)glycinamide that forms hydrogen bound with the OG1 from the highly conserved T262 and NZ from K537 (Fig. 2d). Z1262246195 is a 3,3,3-Tris(fluoranyl)-1-Piperazin that interacts via fluorine atoms 1 with S608 and R617 (Fig. 2e). Fragment Z44584886 is a 4-Fluorophenyl-4-Methyl-Piperazine that is maintained by weak interactions with A264, S293, K389, D540 and L541 (Fig. 2f). Z18618496 is a Methyl n-(Cyclohexanecarbonyl)glycinate that is maintained by weak interactions with L271, D291, P292, S293, M536, K537 and V543 (Fig. 2g). Fragment Z2678251369 is a 2-tert-butyl-5,6,7,8-tetrahydropyrido[4,3-d]pyrimidine that weakly interact with residues A264, S268, S293, L493, M536, D540, L541 and V543 (Fig. 2h). Fragment POB0008 is a Methyl 1-(Methanesulfonyl)-2-Methyl-D-Prolinate that forms hydrogen bond with S268 and weakly interact with residues A264, S268, L271, S293, L493, V533, M536 and K53 (Fig. 2i). Fragment Z3241250482 is a 1,1-bis(oxidanylidene)thietan-3-ol that form hydrogen bounds with K389 and R617, as well as weak interactions with K389, D540, L541, P542, V599, A605, S608, F609 and R617 (Fig. 2j). Fragment POB0128 is a (1s,2r)-2-(Thiophen-3-Yl)cyclopentane-1-Carboxylic acid that form hydrogen bonds with S293 and weakly interacts A264, T267, D291, P292, S293, L493, L541 and V543 (Fig. 2k).

In the structure of PDB 8V7R, two instances of Z56772132, namely 5-methyl-hydantoin, were detected, with the copy designated as chain D occupying the RNA-cleft site. This fragment engages in two hydrogen bonds: one forms between the nitrogen atom (N11) and the oxygen atom of the highly conserved residue D540, measured at 2.8 Å, while the other bond emerges between the oxygen atom (O13) and the NH2 group of R617 at 3.2 Å distance (Fig. 2i). Fragment Z729726784, a 2-cyclopentyl-N-(3-methyl-1,2,4-oxadiazol-5-yl)acetamide, demonstrates interactions with various residues including L271, D291, P292, S293, M536, K537, and V543 (Fig. 2m).

Furthermore, in the structure of ZIKV NS3^Hel^ complexed with Z203039992 (PDB 8UM3), identified as 6-chlorotetrazolo-pyridazine, an intriguing binding pattern emerges. This fragment demonstrates an ability to target the nucleophilic thiol group of C262 by interacting with the main chain nitrogen of R226 and the OE2 atom of G392 via its N1 and N2 atoms (Fig 3a). This ligation of Z203039992 causes a 180° twist of C262 side chain conserved conformation (Fig 3a). It’s noteworthy that C262 represents a highly conserved residue across *Flaviviridae* family, including distant species like HCV (Fig 3c). Positioned within a sub-pocket adjacent to the NTP binding site, C262’s involvement in these interactions could be a unique opportunity for designing novel covalent allosteric or competitive inhibitors targeting *Flaviviridae* helicases.

**Fig. 3.**
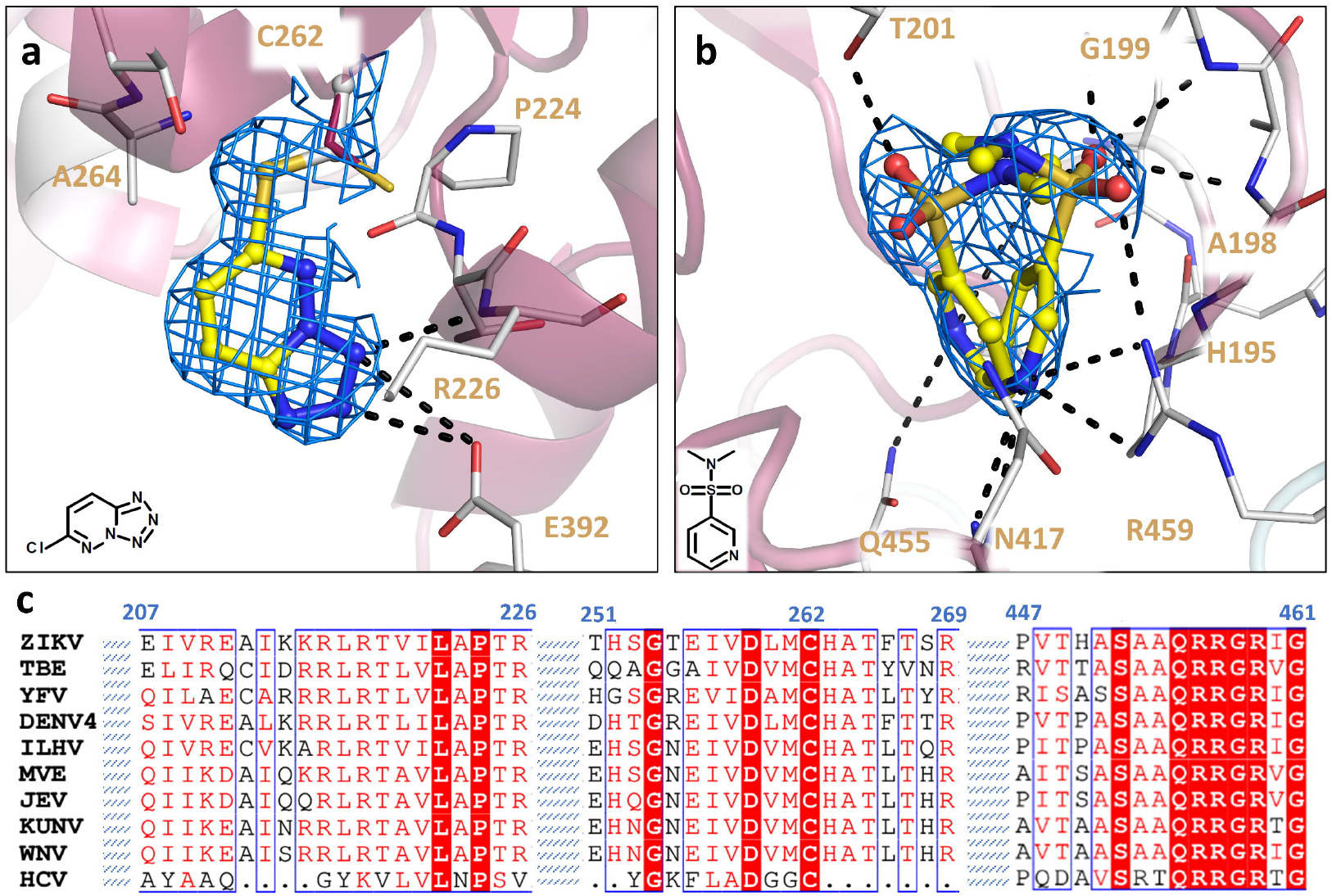
Details of covalent and ATP-binding fragments. (a) Structure of ZIKV NS3^Hel^ in complex with Z203039992. C262 covalently bound is show is depicted with gray carbon, while native apo C262 from PDB 6MH3 is showed in purple (b) Structure of ZIKV NS3^Hel^ in complex with Z751811134 in two overlapping forms. Fragments are shown as sticks with yellow carbons, while protein interacting residues are depicted as gray sticks carbon. Cartoon is colored according to conservancy score calculated with Consurf, where purple indicated more conserved regions and white the less conserved regions. 2Fo-Fc map is depicted as blue mesh, showed here with σ = 1.0. (c) *Flaviviridae* sequence alignment showing selected regions of helicase that are discussed above. Sequence alignment was generated with ESPript and colored with default parameters.

### Fragments for the NTP-binding site

The NTP-binding site represents a highly conserved cavity pivotal for ATP hydrolysis. In line with numerous structures in PDB, our crystal structure of Zika NS3^Hel^ reveals the presence of a phosphate molecule intrinsically occupying the binding site. This molecule likely originates from the expression source and is presumed to compete with any incoming fragment for binding. Therefore, only fragment Z751811134, a N,N-dimethylpyridine-3-sulfonamide was observed in this site. This fragment was found in two mirrored conformations, with the sulfonamide group occupying the position of phosphate (Fig 3b). Fragment Z751811134 interacts with highly conserved residues G199, K200, P196, G197, A198, G199, K200, T201, D285, E286, A317, Q455, R459 and R462, and it’s deposited on PDB under the code 7G9Q. Detailed information including PDB identifiers, fragment names, ligand descriptors, and data collection/refinement statistics can be found in Supplementary Table 1.

### Cluster 3 and surface fragments are crystal artifacts

In Cluster 3 of ZIKV NS3^Hel^, we observed a promiscuous region where we were able to clearly elucidate the structure of 17 fragments, Z198194396, Z126932614, Z425387594, Z1444783243, Z235449082, Z274555794, Z291279160, Z1703168683, Z31222641, Z111782404, Z57821475, Z133716556, Z1619958679, Z1216822028, Z119989094, Z1509711879 and Z85933875, respectively deposited on PDB under the codes 5RHI, 5RHJ, 5RHL, 5RHO, 5RHQ, 5RHS, 5RHT, 5RHU, 5RHW, 7G9L, 7G9N, 7G9O, 7G9P, 7G9Z, 7GA5, 7GA6 and 7GA7. Careful analysis of Cluster 3 indicates that this region is an artifact of crystal packing, and therefore these fragments are too unreliable for seeding a drug design campaign. The same can be said for the 11 surface fragments, Z55222357, Z383202616, Z444860982, Z2048325751, Z56823075, Z1454310449, Z2856434938, Z1324080698, Z385450668, Z758198920 and Z905065822, deposited on PDB under the codes 5RHH, 5RHK, 5RHR, 5RHV, 7G9U, 7G9V, 7G9W, 7GA0, 7GA4, 5RHM, 5RHP, 5RHX, 7G9R, 7G9S and 7G9X. Despite the promising potential of surface fragments in the development of novel protein interaction blockers, our analysis reveals that these fragments are sustained not solely through interaction with the asymmetric unit but also through crystal packing constraints. Consequently, they may not be optimal candidates for design purposes. Detailed information including PDB identifiers, fragment names, ligand descriptors, and data collection/refinement statistics can be found in Supplementary Table 1.

### Resistance analysis

A phylogenetic analysis was conducted on sequencing data from Zika patients, compiled into the NextStrain database by various research groups. This dataset was extracted for further analysis in this study. Our findings indicate that the NS3 Helicase does not exhibit a significantly higher mutation count compared to the rest of the NS3 gene or the broader Zika genome (Fig. 4a).

To visualize these mutation counts, we mapped them onto NS3 Helicase crystal structures. Residue surfaces were colored based on mutation frequency: white for no detected mutations, light red for a single mutation (regardless of recurrence), and darker red for residues with two or more distinct mutation types (e.g., both A1E and A1F, irrespective of their frequency) (Fig. 4b). Focusing on the RNA cleft site, we found that only single mutation types were present in this region (Fig. 4c). Specifically, K537I and R617E were observed four and two times, respectively, while T267S, P292G, S294A, and T390N were each detected once.

## Discussion

After the onset of the COVID-19 pandemic, it became increasingly evident that there is a critical need to prioritize the development of broad-spectrum direct-acting antiviral capable of effectively combating viruses with pandemic potential (24, 25). By applying HTP fragment-screening combined machine learning methods, our prelude group was able to develop a novel oral non-peptidomimetic inhibitor of SARS-CoV-2 Main protease in the open-science format (25–27). Our derived group named AI-driven Structure-enabled Antiviral Platform (ASAP), backed by the NIH Antiviral Drug Discovery (AViDD) program, is committed to employing a similar strategy and ethos to accelerate the development of broad-spectrum, clinic-ready antiviral agents in anticipation of future pandemics, including those caused by flaviviruses.

Despite its promise for therapeutic design, targeting flaviviruses NS3^Hel^ poses significant challenges, particularly due to the scarcity of structural data available for conserved sites. Only a handful of NS3^Hel^ inhibitors, such as benzimidazoles, benzothiazole, pyrrolone, ivermectin, ST-610, and suramin, have been documented in the literature (28–30). However, none of these have been substantiated with comprehensive structural data to support design of new candidates. In the case of HCV, another member of the *Flaviviridae* family, a few structures of NS3^Hel^ in complex with inhibitors have been reported. These include a triphenylmethane series of NTP-site competitors (31) and an indole series that competes with the +4 site in the RNA exit tunnel (32). However, it’s worth noting that ATP competitors can potentially lead to significant toxicological effects, while the pocket accommodating the indole derivatives is wide and groove-like, with inherent flexibility of the protein in this region. Therefore, it’s evident that acquiring new structural data is imperative to facilitate drug design targeting NS3^Hel^.

The ZIKV NS3^Hel^ is segmented into three distinct sub-domains, namely DI, DII, and DIII. This domain exhibits distinctive characteristics with two binding sites: one that selectively engages with single-strand RNA, and the other that binds to a nucleoside triphosphate (NTP) (18). These binding sites are situated within conserved clefts formed between the three NS3^Hel^ subdomains (17, 23, 33). Consequently, these clefts emerge as intriguing areas for exploration in drug design and development (33).

In a prior fragment screening endeavor employing x-ray crystallography on ZIKV NS3^Hel^, a total of 154 fragments were tested, resulting in the detection of only four hits. Notably, all four hits were located at the surface, distantly from both the NTP and the RNA-cleft binding sites or any other conserved region (34). In a comparable study using a thermal-shift assay, researchers screened 500 fragments against DENV helicase but found no hits (35). By applying HTP crystallographic fragment screening, we successfully revealed the structures of 45 fragments at multiple sites for ZIKV NS3^Hel^. Notably, this encompassed the identification of 11 distinct fragments specifically binding to the RNA-cleft site, including a unique covalent hit of the conserved C262 (Fig. 2). A recent publication showed an allosteric inhibitor capable of binding to C441 and C444 on the SARS-CoV-2 helicase (36). The proximity of Z203039992 to both the RNA cleft and NTP site, as well as the other fragments discovered in Cluster 1, suggests its potential in serving as a structural template for the design of novel orthostatic or allosteric inhibitors targeting flaviviruses.

Within Cluster 1, the lower half (Fig. 4c) is bound to regions with lower mutagenic variability, suggesting it may be more suitable for drug discovery. Since elaborated fragments are less likely to target mutable sites, this stability could enhance the likelihood of developing effective therapeutics. In previous studies, we have observed single mutation events occurring at catalytic residues in other proteins, indicating that some low-frequency mutations may result from sequencing errors. Consequently, T267S, P292G, S294A, and T390N should not necessarily be considered less significant than K537I and R617E. Additionally, all bound fragments in the RNA cleft site that interact with residues identified as mutating in this analysis contact sidechains rather than the backbone. Yet, further studies are required to validate the potential of ZIKV NS3^Hel^ as a target for direct acting antivirals.

All together, our findings pave the way for the rapid development of unique inhibitors, specifically targeting the RNA-unwinding activity of flavivirus helicases. Such advancements hold promise for the creation of a new class of direct-acting antivirals.

## MATERIALS AND METHODS

### Cloning, Expression and Purification

The coding region of ZIKV NS3^Hel^ from the MR766 strain (GB: AY632535) was codon optimized for *E. coli* expression and synthesized by Genscript. Region encoding primers CAGGGCGCCATGAGTGGCGCCCTCT-3’ and 5’-GACCCGACGCGGTTACTTTTTCCAGCGG-3’ was amplified and cloned into pETTrx-1a/LIC by Ligation independent cloning method (LIC) (37).

Transformed Rosetta (DE3) *E. coli* (Novagen) cells were grown in ZYM 5052 autoinduction medium (38)supplemented with 50 µg.ml-1 kanamycin and 34 µg.ml-1 chloramphenicol at 37 °C and 200 r.p.m. until OD_600_ of 1.0, followed by 36h shaking at 18 °C. Cells were harvested by centrifugation (4,000 g for 40 min, at 4 ºC) and cell pellet was resuspended in 20 mM Bis-Tris pH 7.0, 500 mM NaCl, 40 mM Imidazole, 10% glycerol supplemented with 4.0 mM dithiothreitol, 2.0 mM phenylmethyl sulfonyl fluoride, 200 µg.L^-1^ lysozyme, 10C□.ml^−1^ benzonase and incubated for 30 min in ice bath. After sonication, cell debris was separated by centrifugation (20,000 g, 20 min, 4°C) and clarified material was loaded into HisTrap HP 5.0 mLpre-equilibrated with 20 mM Bis-Tris pH 7.0, 500 mM NaCl, 40 mM Imidazole, 10% glycerol. After washing, protein was eluted with a gradient of the same buffer containing 500 mM imidazole. Protein 6xHis-Thioredoxin tag was cleaved with TEV protease during O.N. dialysis in 20 mM Bis-Tris pH 7.0, 500 mM NaCl, 40 mM Imidazole, 10% glycerol supplemented with 4.0 mM dithiothreitol. A second passage through HisTrap HP 5.0 mL was used to remove cleaved tag and TEV. Protein was concentrated and inject into size-exclusion chromatography on a Superdex 16/60 75 column (GE Healthcare) pre-equilibrated in buffer 20 mM Bis-Tris pH 7.0, 500 mM NaCl, 10% glycerol for polishing. For crystallization, samples where concentrated to 5.0 mg.ml^-1^ based on theoretical extinction coefficient of 64,775 M^-1^.cm^-1^ at 280 nm as reference (39).

### Crystal growing and optimization for screening

Initial crystals were grow in a crystallization solution derived from Morpheus HT Crystallization Screen (Molecular Dimensions), comprising 0.12 M NPS Mix (consisting of 0.3 M Sodium phosphate dibasic dihydrate, 0.3 M Ammonium sulphate, and 0.3 M Sodium nitrate from Molecular Dimensions), along with 0.1 M MES/Imidazole at pH 6.5 (Molecular Dimensions), and 33% Precipitant Mix 4 (11% MPD, 11% PEG 1,000, and 11% PEG 3,350 from Sigma Aldrich), maintained at 20°C. An initial model was obtained at a resolution of 1.9 Å using a Rigaku Micromax-007 HF X-ray diffractometer, processed within the *P*2_1_ space group using XDS and Aimless (40, 41), and subsequently solved through molecular replacement employing the PDB entry 5GJB as a reference model with Phaser (42) and refined with phenix.refine (43). The resulting structure was deposited under the code 6MH3 (Supplementary Table 1).

For fragment screening, crystals were treated by aspirating and vortexed them with 100 µL of the crystallization solution along with glass beads. Subsequently, 0.05 µL of seed stock was introduced into each drop, maintaining a 1:1 ratio of protein to crystallization solution. This process led to the acquisition of viable crystals in approximately 60-70% of the drops within 48 hours of the plate setup. To determine solvent tolerability, crystals were incubated with dimethyl sulfoxide (DMSO) in concentrations of 10-30% and incubated for periods of 1-3h at room temperature.

### Fragment screening and data collection and processing and analysis

The fragment screening was performed using the XCHEM infrastructure available at Diamond Light Source. The soaking was done by transferring fragments from DSi-Poised Library (44), Minifrags (45), Fraglites (46), Peplites (47), York 3D (48), SpotXplorer (49), and Essential Fragment Library (Enamine) to the crystal drop using ECHO Liquid Handler at proportion of 20%. Crystals were incubated for 2h at room temperature and harvested using Crystal Shifter (Oxford Lab Technologies), mounted on loops and flash frozen in liquid nitrogen.

All data was collected at I04-1 beamline at Diamond Light Source at 100 K and processed with automated pipelines using combined software’s, with resolution cutoff defined algorithmically (40, 50–52). All further analysis was performed using XChemExplorer (53). Initial density maps were generated using DIMPLE (52) and PDB 6MH3 as template model. Ligand restrains were generated with ACEDRG and GRADE (54, 55). A ground state model was constructed utilizing PanDDA by amalgamating 100 apo datasets (56). This model, deposited under the code 5RHY (Supplementary Table 1), served as the template for molecular replacement for the full analysis. Event maps were calculated with PanDDA (56), and ligands were modeled using Coot (57) Models were refined with Refmac (58), Buster (54) and phenix.refine (43), and models and quality annotations cross-reviewed. Coordinates, structure factors and PanDDA event maps for all data sets are deposited in the Protein Data Bank (PDB). Data collection and refinement statistics, PBD codes and ligand identifier are available in Supplementary Table 1. Figures were made using PyMOL (Schrödinger), ChimeraX (59), Consurf (60) and E/ENDScript (61). Protein alignment in Fig. 1b used PDBs 5JPS, 5Y4Z, 6ADW, 6ADX, 6ADY, 6MH3, 5VI7, 5GJB, 5GJC, 5JMT, 5JRZ, 5JWH, 5K8I, 5K8L, 5K8T, 5K8U, 5TXG, 5Y6M, 5Y6N, 7V2Z, 8CXG, 8CXH, 8CXI, 5MFX, 6RWZ, 6S0J, 7WD4, 2BHR, 2BMF, 8GZQ, 8GZR, 2JLS, 2VBC, 2WHX, 2WZQ, 2JLR, 2JLU, 2JLV, 2JLW, 2JLX, 2JLZ, 5XC6, 5YVJ, 5YVU, 5YVV, 5YVW, 5YVY, 5YW1, 2JLQ, 2JLY, 5XC7, 7XT0, 2V8O, 2WV9, 2QEQ, 2Z83, 8JIX, 2V6I, 2V6J, 1YKS, 5FFM, 7BLV, 7BM0, 7NXU, 7OJ4, 7JNO, 7V4Q, 7V4R, 6M40, 1HEI, 4OJQ, 4OK3, 4OK5, 4OK6, 4OKS, 4WXR, 1A1V, 5GVU, 5WSO, 3KQH, 3KQK, 3KQL, 3KQN, 3KQU, 4CBL, 5E4F, 2ZJO, 4CBG, 4CBH, 4CBI, 5MZ4, 5WX1, 1CU1, 2F55, 3O8B, 3O8C, 3O8D, 3O8R, 3RVB, 4A92, 4B6E, 4B6F, 4B71, 4B73, 4B74, 4B75, 4B76, 4CBM, 4WXP, 5FPS, 5FPT, 5FPY, 8OHM and 5WDX.

### Resistance analysis

Zika patient sequencing data was analyzed using NextStrain <u>PubMed</u>, with phylogenetic trees generated from the retrieved data. The NextStrain API was queried via GET requests, and the resulting JSON files were processed to extract mutation counts per residue. Mutations were quantified by counting the number of events at the tips of the phylogenetic tree.

Additionally, the workflow includes an entropy-based analysis, though this approach was not utilized in this manuscript. Each tree leaf was traced back to its original residue to determine the wild-type sequence. The number of unique mutants per residue was then calculated, and surface representations of Zika NS3Hel were visualized in PyMOL, where residues with zero mutations were colored white, and those with increasing mutation diversity were shaded progressively red.

For this analysis (starting from the NextStrain query) an MIT-licensed software product called Choppa was built [https://zenodo.org/records/11100680] that can generate similar interactive representations on comparable datasets such as the ones derived from Deep Mutational Scanning data [https://pmc.ncbi.nlm.nih.gov/articles/PMC4410700/].

## Supporting information

crystallography table

## Data availability

All detailed protocols and associated data can be accessed at https://asapdiscovery.org/. Crystallographic coordinates and structure factors for all structures have been deposited in the PDB with the following accession codes 6MH3, 5RHG, 7G9K, 7G9M, 7G9T, 7G9Y, 7GA1, 7GA2, 7GA3, 8UM3, 8V7R, 8V7U, 7G9Q, 5RHI, 5RHJ, 5RHL, 5RHO, 5RHQ, 5RHS, 5RHT, 5RHU, 5RHW, 7G9L, 7G9N, 7G9O, 7G9P, 7G9Z, 7GA5, 7GA6, 7GA7, 5RHH, 5RHK, 5RHR, 5RHV, 7G9U, 7G9V, 7G9W, 7GA0, 7GA4, 5RHY, 5RHM, 5RHP, 5RHX, 7G9R, 7G9S and 7G9X.

## Acknowledgements

Research reported in this publication was supported in part by NIAID of the National Institute of Health under award number U19AI171399. The content is solely the responsibility of the authors and does not necessarily represent the official views of the National Institute of Health. GO and ASG acknowledge Fundação de Amparo à Pesquisa do Estado de São Paulo (FAPESP project 2013/07600-3). Authors acknowledge Diamond Light Source access to I04-1 beamline and XCHEM facilities through proposals lb20289 and lb32627.

## Notes

### Competing Interest Statement

ASG consults for DNDi and MMV

### Summary of Updates

Two authors added from last version

